# Mature sperm small RNA profile in the sparrow: implications for transgenerational effects of age on fitness

**DOI:** 10.1101/520759

**Authors:** Wayo Matsushima, Kristiana Brink, Julia Schroeder, Eric A. Miska, Katharina Gapp

**Affiliations:** Gurdon Institute, University of Cambridge, Tennis Court Rd, Cambridge, CB2 1QN, UK.; Wellcome Trust Sanger Institute, Wellcome Genome Campus, Hinxton, CB10 1SA, UK.; Department of Genetics, University of Cambridge, Downing Street, Cambridge, CB2 3EH, UK.; Department of Life Sciences, Imperial College London, Silwood Park Campus, Ascot, SL5 7PY, UK.

## Abstract

Mammalian sperm RNA has recently received a lot of interest due to its involvement in epigenetic germline inheritance. Studies of epigenetic germline inheritance have shown that environmental exposures can induce effects in the offspring without altering the DNA sequence of germ cells. Most mechanistic studies were conducted in laboratory rodents and C.elegans while observational studies confirm the phenotypic phenomenon in wild populations of humans and other species including birds. Prominently, paternal age in house sparrows affects offspring fitness, yet the mechanism is unknown. This study provides a first reference of house sparrow sperm small RNA as an attempt to uncover their role in the transmission of the effects of paternal age on the offspring. In this small scale pilot, we found no statistically significant differences between miRNA and tRNA fragments in aged and prime sparrow sperm. These results indicate a role of other epigenetic information carriers, such as distinct RNA classes, RNA modifications, DNA methylation and retained histones, and a clear necessity of future studies in wild populations.

## Introduction

Previously, sperm cells were thought to contribute little but the paternal haplotype to the zygote(1). Along the information provided by DNA sequences sperm cells carry potential epigenetic information in form of modifications of the DNA such as DNA methylation and post translational modifications of histones, the proteins that DNA is wrapped around. Yet DNA methylation is reprogrammed shortly after fertilization in the zygote and most histones are replaced by protamins during sperm maturation(2). The discovery of complex RNA populations in sperm provides an alternative reprogramming independent epigenetic information vector(3, 4). In addition to relatively low abundant messenger RNA (mRNA), sperm cells have been found to contain small non-coding RNA (ncRNA) such as microRNA (miRNA), Piwi-interacting RNA (piRNA) and transfer RNA (tRNA) fragments(3, 5, 6). Whilst the function for most sperm RNA remains unknown, there is evidence that sperm miRNA contribute to early embryonic development(7, 8), suggesting that sperm RNA has fitness consequences on both male and female offspring. A number of studies have also shown that sperm non coding and coding RNA can change in response to life experiences(9–17). This opens the intriguing possibility that alterations in sperm RNA may relay information from father to offspring, potentially about the environment. Indeed, the capacity of altered sperm RNA to transmit information from one generation to the next has been shown by our own work and that of others(9, 15)(10)(11)(12). The effects of a variety of environmental conditions on the offspring was recapitulated using sperm RNA injection of exposed males into naïve fertilized oocytes. These findings provide conclusive evidence that one functional vector for environmentally induced paternal epigenetic germline inheritance are sperm RNA(18–21). As sperm cells contain distinct populations of RNAs there has been great interest in understanding which sub population is functionally active(6). Most studies so far focused on the role of small RNA including miRNAs(12, 14, 21), tRNA fragments(11) and their modifications(10, 22) while our most recent study also implicates long RNA in sperm as an active intergenerational information carrier(15). Importantly, these studies were conducted in laboratory mice in highly controlled environments. Mechanistic studies in other species are lacking and thus the relevance of sperm RNA for real life non-genetic phenomena remains to be demonstrated.

One example of a non-genetic, intergenerational effect is paternal age on offspring fitness, often referred to as the Lansing effect. The Lansing effect describes decreased offspring lifetime fitness with increased paternal age. Previous studies have suggested an epigenetic underpinning of this effect(23, 24). A Lansing effect has been identified in a diverse range of taxa, including birds(23, 25–28). The transgenerational consequences of the Lansing effect can influence a population’s rate of genetic and phenotypic change, which in turn affects the population’s ability to adapt and persist(29). Yet, despite its societal and ecological importance for our understanding of the evolution of ageing and longevity, the mechanistic underpinnings of the Lansing effect are still unknown(23, 24, 30, 31).

The Lansing effect has previously been demonstrated in wild house sparrows (*Passer domesticus*), small passerines considered an important model species in ecology and evolution (32). Indeed, in a natural population, we found that aged sparrows sired offspring with lower life-time fitness(24). While previously, molecular ecological research in this species has been limited to a small number of microsatellite markers and sequencing of a few immune genes(33–37), more recently, a genome assembly and transcriptome sequencing annotation from sparrow somatic tissue has been reported(38, 39) yet the sparrow small non-coding RNome to date is unknown. The implication of comparative expression studies hence have thus far been hindered by a lack of readily available conventional analysis tools such as entries in the miRbase, a database for known miRNA annotations(40). Thus, to facilitate future analysis of sperm RNA in epigenetic transgenerational inheritance a reference set of known house sparrow miRNAs is essential.

Here we present the first isolation and characterization of house sparrow sperm sncRNA with a focus on miRNAs and tRNA fragments. We then describe a first attempt to test for differences in small RNA payload between prime and aged male house sparrows. The identified miRNAs provide a valuable tool for upcoming molecular analysis of gene expression regulation in this interesting model system. Hence, this study sets the ground for future mechanistic research on the Lansing effects in sparrows and therefore has implications for our understanding of the evolution of aging and reproductive lifespan.

## Methods

### Animals

Captive house sparrows, *Passer domesticus*, (n=209) were kept at Silwood Park, Imperial College London in seven aviaries, each holding reproductively active sparrows of mixed sex and age. We identified individuals by uniquely numbered and colored leg rings(41). The age of each individual, and the genetic parentage was available for all individuals(41). Sparrows were between 1-13 years old, and breeding group compositions were chosen to avoid inbreeding (for more details, see (41)). We provided and food *ad libitum*. Sparrows breed seasonally in spring and summer, and undergo dramatic gonadal development in response to length of photoperiod and air temperature(42). Therefore, samples were collected during the breeding season, between April – June 2018 when sperm production in males is at its highest(43). 4 CBLT mice were cohoused in one cage in a temperature and humidity-controlled facility under a non-reversed light-dark cycle, and food and water were provided *ad libitum*.

### Experimental procedure/ Sample collection

We classified male sparrows as either ‘aged’, 8-13 years old (n=9) or as ‘prime’, 1-4 years young (n=8), because in the wild, sparrows increase their reproductive output early in life and plateau at four years of age, after which reproductive senescence sets in(44). Reproductively active sparrows store sperm at the terminal end of the vas deferens. This storage results in a swelling, the cloacal protuberance. Sperm samples were obtained by cloacal massage(45). We massaged the cloacal protuberance gently until ejaculation occurred typically near immediately, and then we collected the ejaculate with a glass capillary.

Male mice of 2.5 months of age were sacrificed by cervical dislocation during their inactive cycle. We dissected their cauda epididymis and vas deferens and placed them in M2 medium. Mature sperm cells were separated from potential somatic contamination by swim-up(46) followed by somatic lysis(11).

### Sparrow sperm purification

We purified samples by somatic lysis: samples were washed with 500 *μ*l of M2 medium (SIGMA), followed by centrifugation at 3000 *g* for 3 minutes. The supernatant was discarded and the pellet was resuspended in 500 *μ*l of somatic lysis buffer and placed on ice for 10 minutes before repeating the centrifugation step. We then discarded the supernatant and added 500 *μ*l of PBS buffer to wash the sample before a final centrifugation step. The final supernatant was discarded and the sample frozen at minus 80°C.

### RNA extraction and quality control

Total RNA was extracted from purified sperm using a slight modification of a standard Trizol protocol (Thermo Fisher Scientific). In brief, sperm samples were resuspended in 1000 *μ*l TriSURE (Bioline) and further homogenised by passing through a syringe (26G) 2-3 times. Subsequently, 200 *μ*l of chloroform (Hi-Media) was added and was mixed vigorously. After 2-3 minutes the samples were then centrifuged for phase separation. The aqueous phase was then retrieved and mixed with an equal amount of chloroform. Again, the upper aqueous phase was recovered after centrifugation. For precipitation equal volume of Isopropanol (Hi-Media) and 2 *μ*l of glycogen (20mg/ul) were added to the aqueous phase. The RNA pellet was washed twice with 75% ethanol (Hi-Media). The pellet was kept to air dry and resuspended in 10 *μ*l nuclease free water. miRNA quantity was assessed using a Qubit fluorometer (microRNA Assay, Life Technologies). RNA purity was determined by Agilent 2100 Bioanalyser (RNA 6000 Pico Kit, Agilent Technologies).

### Library preparation

Total RNA samples were used to create small RNA libraries for Next Generation Sequencing (NGS). Small RNA libraries were generated using the TruSeq Small RNA Sample Prep Kit (Illumina) with slight modification to the manufacturer’s instructions. An average of 29 ng of sparrow sperm total RNA and 20 ng of mouse sperm RNA was used for library preparation according to the manufacturer’s recommendations with the following modifications: 3′ and 5′ adapters were diluted in a 1:4 ratio with RNase free water and PCR amplification was increased to 18 cycles. The concentration of the libraries of cDNA was analysed using Qubit 2.0 Fluorometer (dsDNA HS Assay), followed size assessment using Agilent 2200 TapeStation (HS D1000 ScreenTape, Agilent Technologies). Sequencing was performed on an Illumina HiSeq2500 with 50-bp single-end read length.

### Sequencing analysis

We carried out a quality analysis of the reads before and after adapter trimming using FastQC v.0.11.2(47). After adapter removal using CutAdapt v.1.7 (48) we discarded the reads with inserts of <18-bp, and those where no adapters were found. For tRNA analysis, “CCA” sequence at the 3’ end of reads was further removed since this trinucleotide is non-templatedly added during the tRNA maturation process. Since the house sparrow genome is available(39), but miRNA entries are lacking we used two modalities for further analysis:

1. Using zebra finch as a reference genome We mapped the trimmed reads to the reference genome of *Taeniopygia guttata* (Ensembl, taeGut3.2.4) using the STAR aligner(49) with following parameters: “-- outFilterMultimapNmax 5000 --winAnchorMultimapNmax 5000 -- outFilterMismatchNoverLmax 0.05 --outFilterMatchNmin 16 –outFilterScoreMinOverLread 0 --outFilterMatchNminOverLread 0 --alignIntronMax 1 --alignEndsType EndToEnd -- outFilterType BySJout”. The number of counts mapping to entries of miRBase v.22(40) and/or GtRNAdb tRNA annotations (teaGut324)(50, 51) was determined using FeatureCounts(52). The mapped reads were then analysed for differential expression between the two age classes using DESeq2 workflow v.1.11.22(53). Annotations with no, or single read counts, across all samples were removed. Male sparrows in their prime were set as the reference level for all analysis. Multiple comparison adjustment was carried out, using the Benjamini-Hochberg (BH) adjustment to account for false positives resulting from multiple testing.
2. Using mirDeep2 Reads were mapped to *Passer domesticus* genome (v.1.0)(54) with the mapper.pl script from mirDeep2 and the potential miRNA-coding regions were predicted by mirDeep2.pl script. To improve its prediction accuracy, mature miRNA sequences of zebra finch (miRBase v.22) were provided in this prediction step. The number of reads assigned to each predicted miRNA was calculated by quantifier.pl script and then analysed by DESeq2 as above.

Comparison with mouse sperm Mouse sperm small RNA sequencing reads were trimmed as described above and aligned to the mouse genome (mm10) using the STAR aligner(49) with following parameters: “-- outFilterMultimapNmax 5000 --winAnchorMultimapNmax 5000 --outFilterMismatchNmax 0 --alignIntronMax 1 --alignEndsType EndToEnd”. Downstream analyses were performed in the same way as in the sparrow dataset under modality (1).

## Results

### Somatic lysis on samples obtained by cloacal message yields pure sperm

To evaluate the possibility to yield a pure cell population of mature sperm, we subjected samples obtained by cloacal massage to somatic lysis (Figure 1a) before extracting RNA using the Trizol method, and examined the RNA profile on the bioanalyzer for ribosomal peaks. Prominent ribosomal peaks are indicative of somatic cell contamination, since maturing sperm undergoes ribosomal cleavage(55). We observe a peak in the region below 200 nucleotides (nt) typical for small non coding RNA but no peaks in the size range of ribosomal RNA (Figure 1b, Supplementary figure 1a,b), suggesting a pure mature cell population.

**Figure 1:**
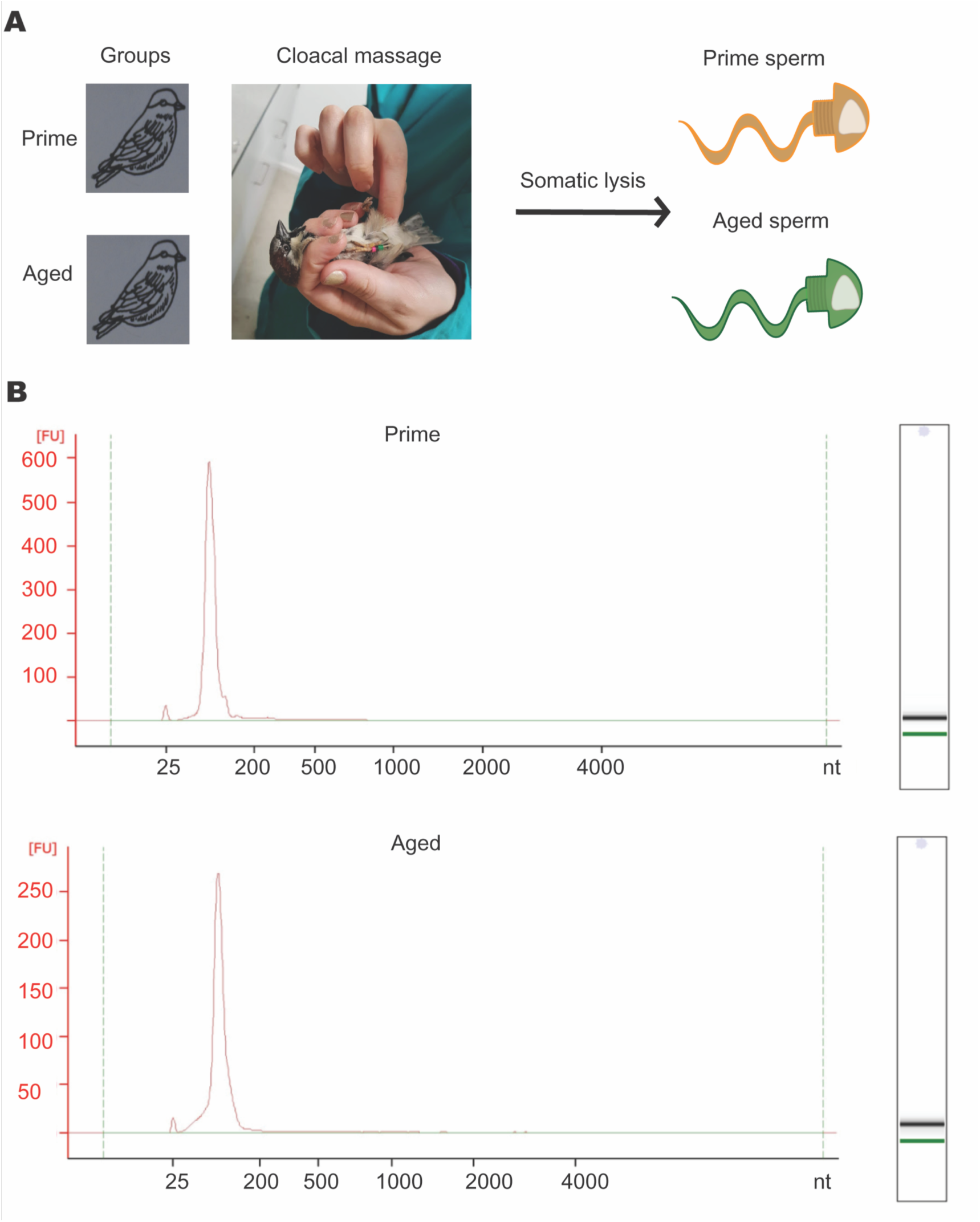
(a) Experimental design (b) Total RNA bioanalyzer Profiles of prime and aged sparrow sperm total RNA.

### Small RNA payload of sperm depicts RNA species of different sizes and classes

Using next generation sequencing we produced small RNA libraries of the sparrow sperm RNA and investigated their nucleotide composition and RNA class identity. Collapsing all reads from aged and prime sperm RNA libraries respectively revealed a distinctive size profile between 18 and 50 nt (Figure 2a,b). Both age groups show a prominent peak at 32 nt, corresponding to the size of tRNA fragments, but potentially also piRNAs. piRNAs can be identified by uracil in the first position, yet such bias is not observed. A second less pronounced peak is observed at 22 nt in prime sperm RNA libraries, which is indicative of miRNAs. Samples of aged sparrows show higher peaks at 18 nt. When counting reads with the same sequence once only, no peak is observed at 32 nt (Figure 2c,d). The inspection of the relative abundance of reads mapping to distinct RNA subclasses showed overall the highest abundance for tRNA mapping reads, followed by rRNA mapping reads, miRNA mapping reads and reads mapping to mitochondrial RNA among others (Figure 2e).

**Figure 2:**
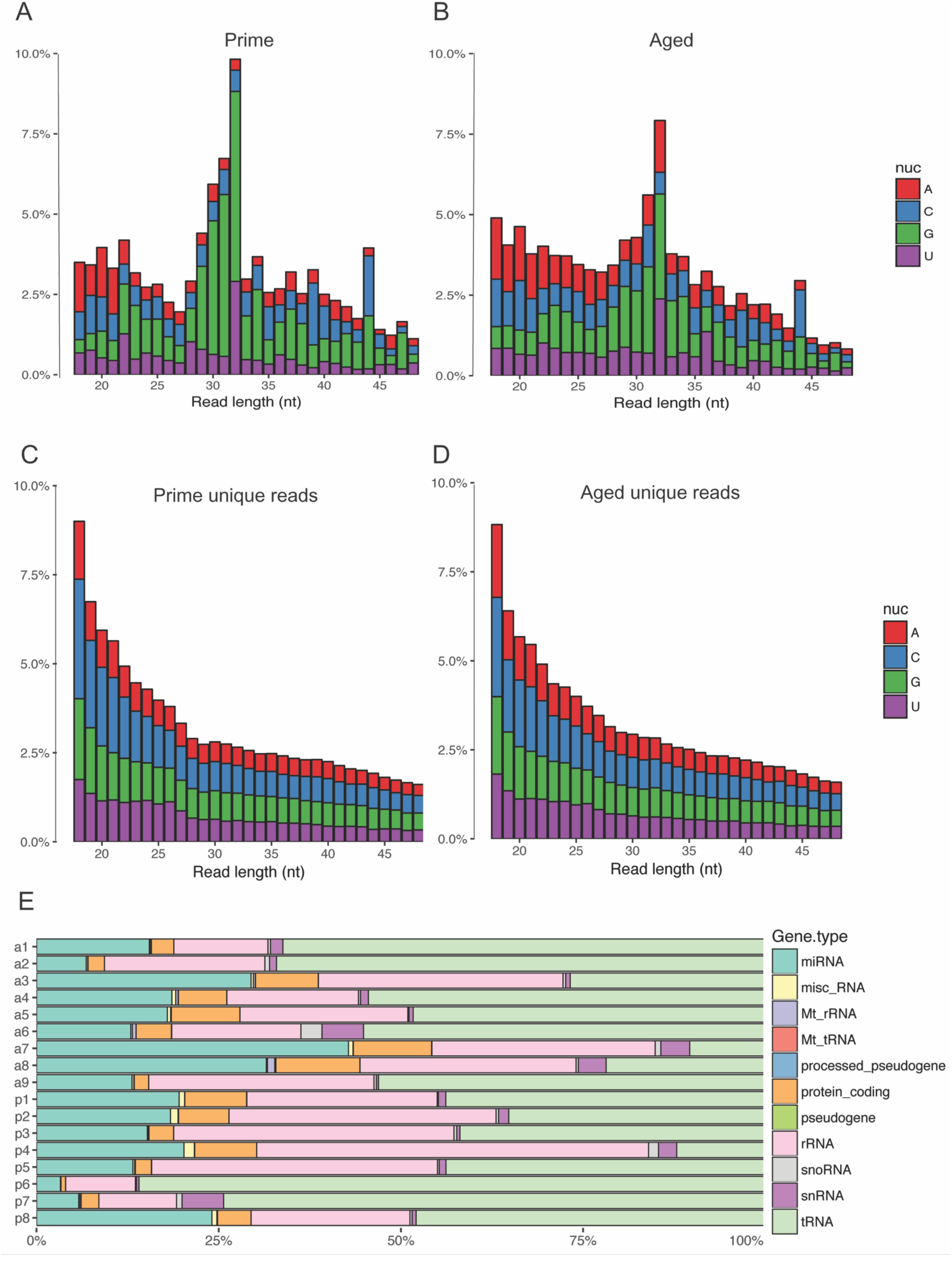
Size distribution of small RNA next generation sequencing reads from prime and aged sparrow sperm using (a,b) representing all reads (c,d) representing unique reads only. Colours depict first base identity. (e) Relative abundance of reads mapping to different RNA classes in each sample.

### Analysis of sparrow sperm small RNA sequencing indicates abundant presence of miRNAs

The mapping of sequencing reads to the zebra finch genome and the respective mirBase entries identified the presence of 334 miRNAs in sparrow sperm (Supplementary Figure 2). A comparison of the 10 most abundant sparrow sperm RNA sequencing reads mapping to zebra finch miRNAs and mouse sperm RNA sequencing reads mapping to mouse miRNAs revealed a conserved high abundance of tgu-miR-let-7a, tgu-miR-let-7f and tgu-miR-10a-5p (Figure 3a,b; Supplementary table 1).

**Figure 3:**
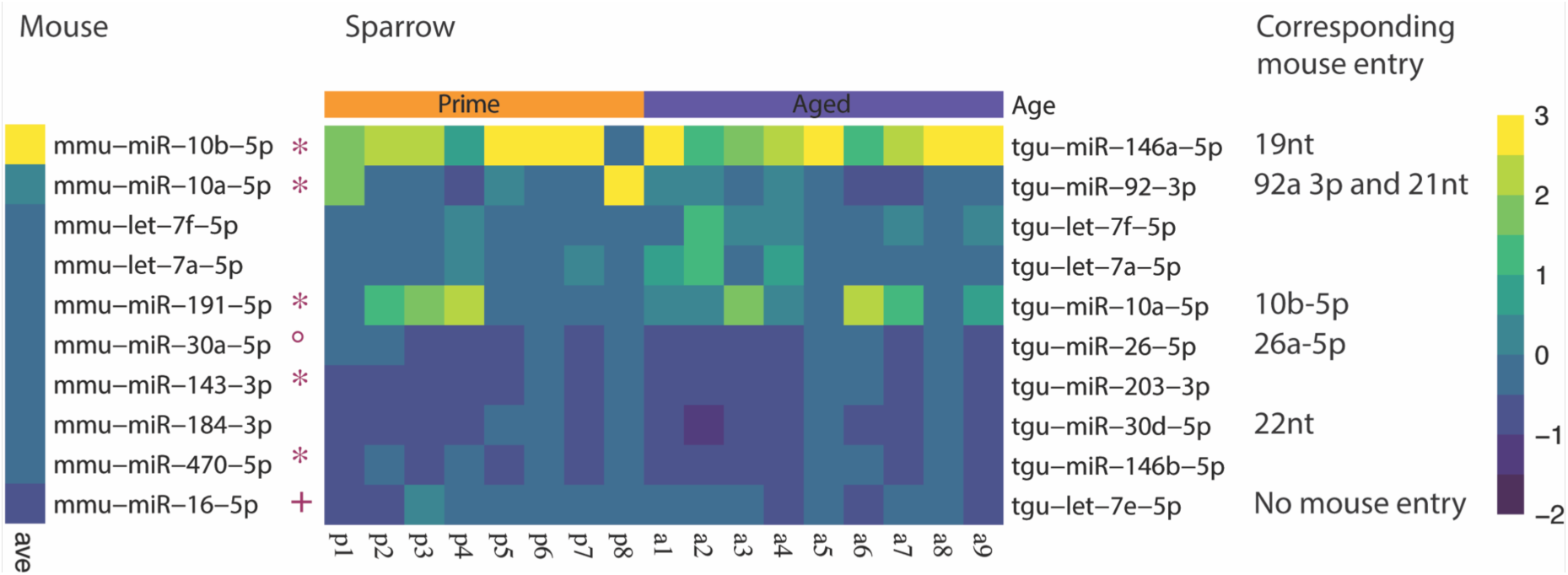
Heatmap illustrating 10 most abundant miRNAs in (a) mouse and (b) sparrow sperm as determined by quantifying the number of sequencing reads mapped to zebra finch miRNA annotations in miRBase. Mouse miRNAs marked with * do not have respective miRNA entries in zebra finch miRBase. Mouse miRNA marked with ° is identical with zebra finch tgu-miR30b-5p. Mouse miRNA marked with + is identical to zebra finch tgu-miR-16-5p apart from 1 mismatch. Colour code represents a z-score.

### miRDeep2 detection of miRNAs in sparrow sperm identifies novel and conserved miRNAs

Since the sparrow miRNA sequences might deviate from the zebra finch miRNA sequences, we explored an alternative analysis of sparrow small RNA based on *de novo* discovery of miRNA annotation by the miRDeep2 algorithm(56). This approach allows unbiased detection of sparrow miRNAs independent of zebra finch miRNA annotation. mirDeep2 depicted a broad range of potential miRNAs in the sparrow sperm small RNA sequencing dataset (Figure 4a, Supplementary table 3), some of which share seed sequences with known zebra finch miRNAs, but some of which do not (Figure 4b, Supplementary table 4).

**Figure 4:**
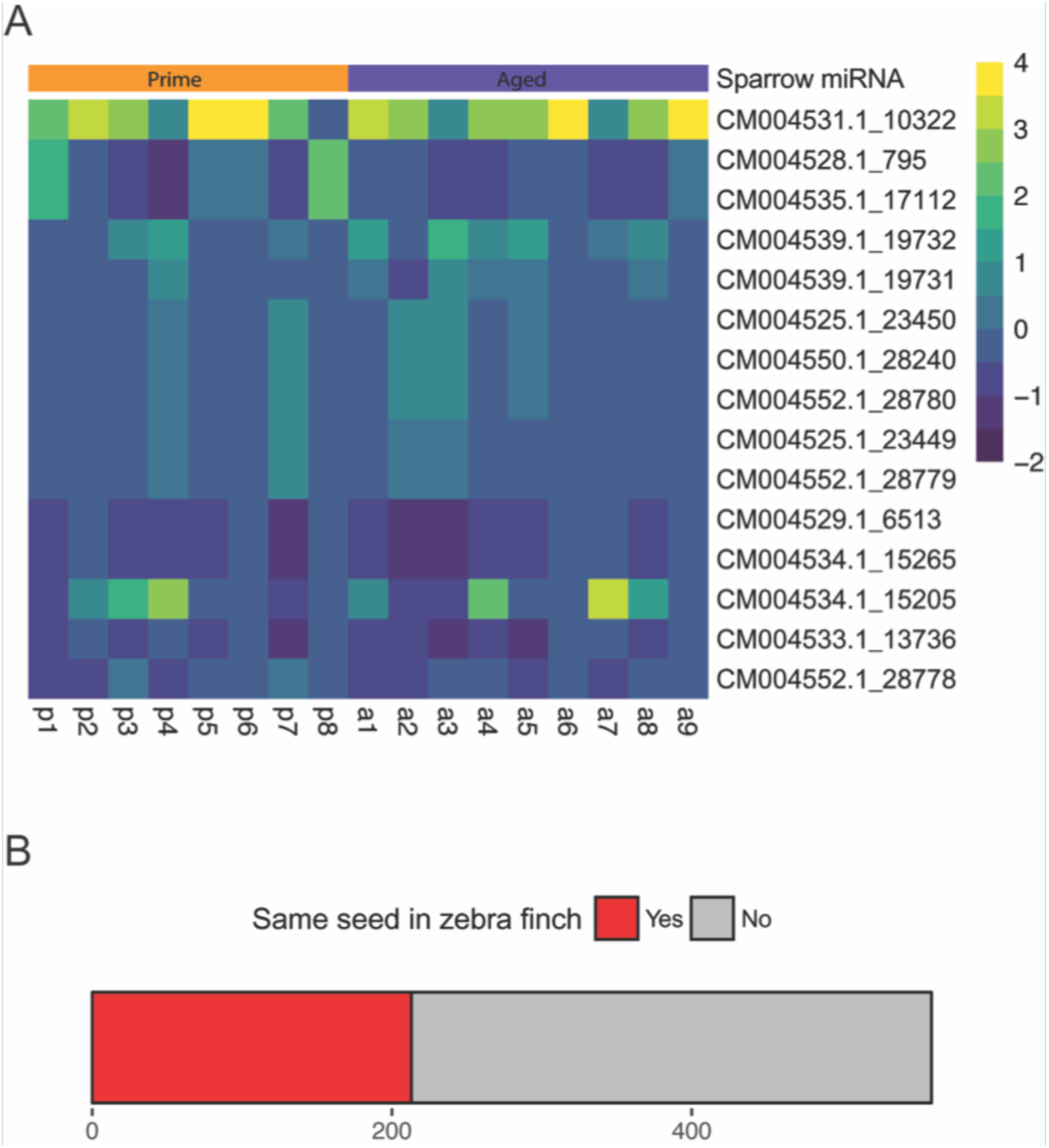
(a) Heatmap illustrating 10 most abundant miRNAs in sparrow sperm as determined by miRDeep2 miRNA de novo analysis. Colour code represents a z-score. (b) Relative amount of shared seed sequence between miRNAs identified by miRDeep2 and known zebra finch miRNAs.

### Differential miRNA expression analysis reveals no significant differences between prime and aged sparrow sperm samples

To investigate the potential impact of sparrow age we used the DESeq2 package to test for the differential transcript abundance in miRNAs identified with zebra finch miRBase as a reference and those identified with miRDeep2. Neither analysis revealed significant difference in miRNA abundance between aged and sparrows in their prime (Figure 5a,b, Supplementary table 4a,b).

**Figure 5:**
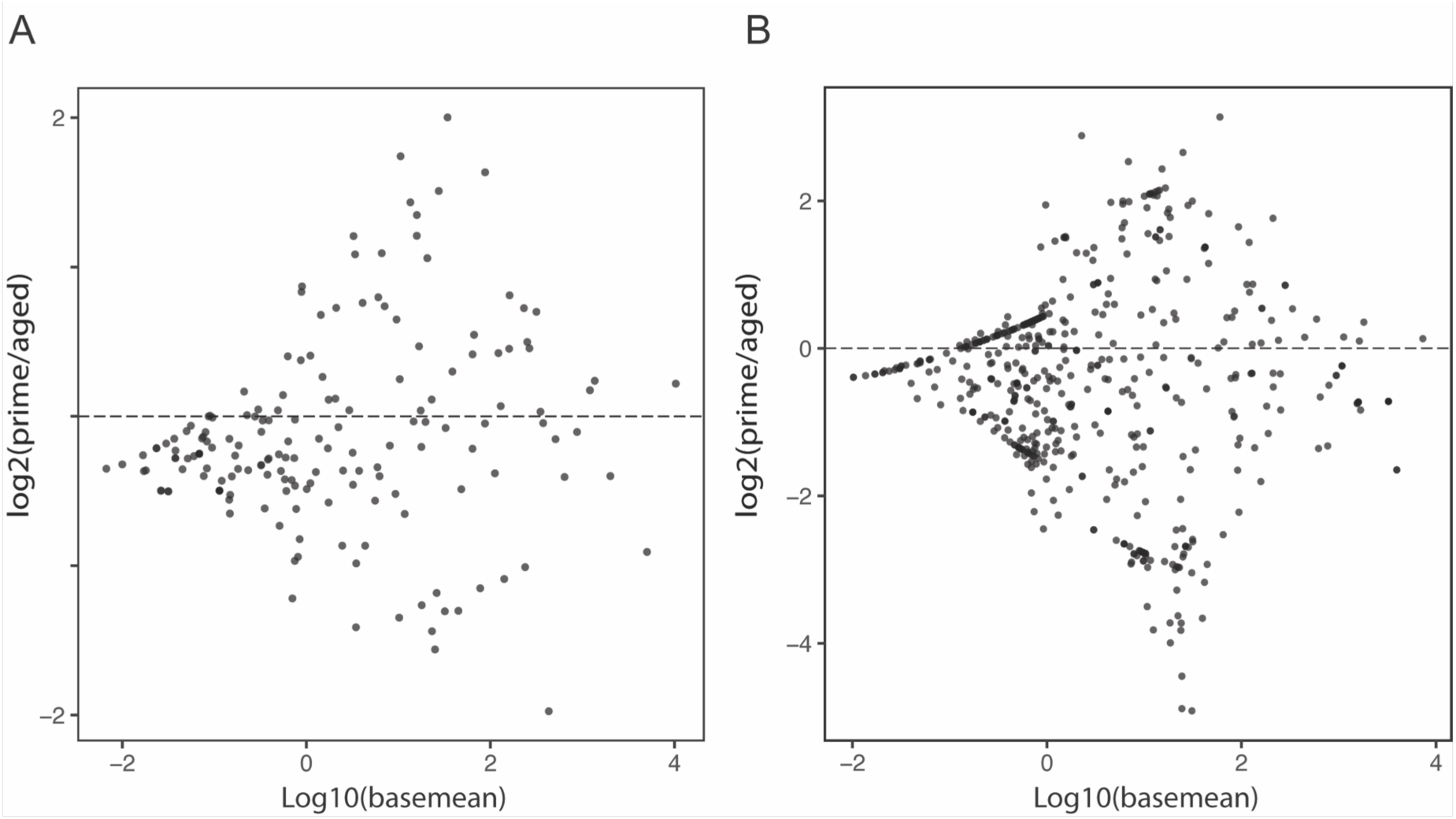
MA plot depicting results of DESeq2 differential miRNA expression analysis (a) using the zebra finch miRBase entries and (b) using miRDeep2.

### Analysis of sparrow sperm small RNA sequencing data indicates abundant presence of reads mapping to tRNA fragments

To further study the identity of sparrow sperm RNA we also quantified the sequencing reads mapped to tRNA annotated regions of the zebra finch genome. We detected a variety of reads mapping to tRNA fragments (Figure 6a, Supplementary table 5a) in aged and prime sparrow sperm libraries. To further investigate potential differences in their abundance depending on sparrow age we performed a differential expression analysis using DESeq2. This analysis did not detect any statistically significant differential abundance of tRNA fragments (Figure 6b, Supplementary table 5b).

**Figure 6:**
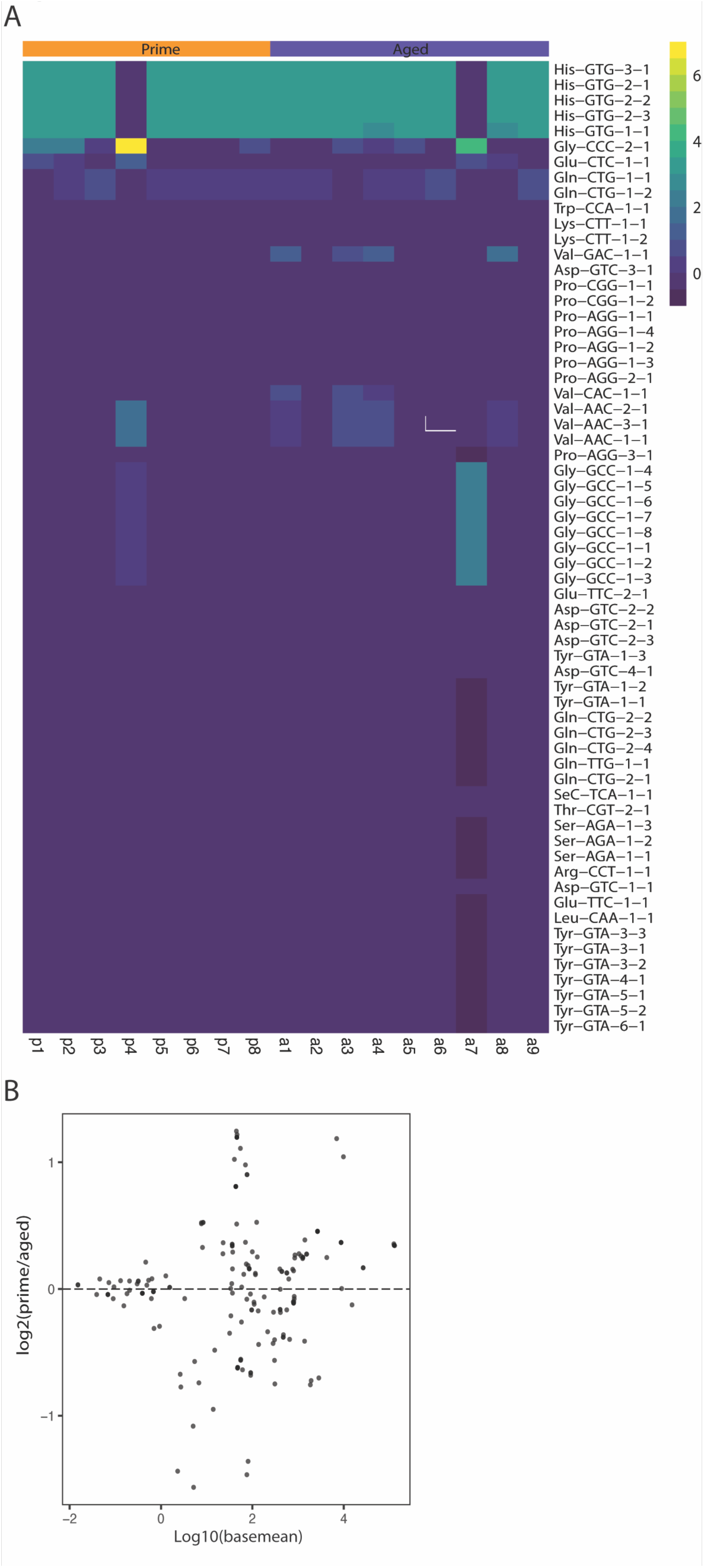
(a) Heatmap illustrating 10 most abundant tRNA fragments in sparrow sperm as determined by quantifying the number of sequencing reads mapped to zebra finch tRNA annotations. Colour code represents a z-score. (b)MA plot depicting results of DESeq2 differential miRNA expression analysis using the zebra finch miRBase entries.

## Discussion

The current study provides the first analysis of sperm small RNA in the house sparrow, with the long-term goal to enable comparative studies of environmentally exposed sperm and its contribution to epigenetic germline inheritance.

We isolated a pure sperm population from sparrow cloacal sperm samples, as assessed by the RNA size profile obtained from the bioanalyzer (Figure 1, Supplementary figure 1b). The absence of peaks of the size of ribosomal RNA, indicative of somatic, actively translating cells, excludes contamination. This is essential since a minute contamination of somatic cells would drastically influence our small RNA readout due to the high abundance of RNA in somatic cells as compared to mature sperm. Interestingly, the bioanalyzer profile also revealed a peak of small RNA at 60 nt (Supplementary figure 1a), not observed in mouse sperm (8). This peak could potentially indicate the presence of an additional RNA class of small nucleolar RNA, involved in the regulation of RNA modifications(57), yet the identity of this peak of sparrow sperm RNA remains to be determined.

The pure population of sparrow sperm yielded enough RNA (a minimum of 20ng/individual) to proceed to small RNA library generation for next generation sequencing. The obtained data were analyzed using the zebra finch, a phylogenetically closely related relative of the sparrow, as a reference genome, as to obtain miRNA annotations available from miRBase.

The small RNA library size profile of both aged and prime sparrow sperm samples demonstrated peaks at 32 nt, indicative of tRNA fragments or piRNAs (Figure 2a,b). Identical tRNA fragments and piRNAs are encoded at multiple genomic loci resulting in a higher prevalence of identical reads and ultimately in more overall reads of these classes. The disappearance of the 32 nucleotide peak observed in the size analysis counting unique reads only (Figure 2c,d) indeed suggests 32 nucleotide long RNA to consist of a less varied set of sequences such is tRNA fragments or piRNAs. Sperm of sparrows in their prime also had a small peak at 22 nt which corresponds to the size of miRNAs (Figure 2a). While this peak is absent in aged sperm, this group shows an additional peak of 19 nt, potentially also corresponding to shorter miRNAs (Figure 2b).

The potential size difference between prime and aged sperm at 22 nt does further not persist when examining unique reads only, indicating no generic difference in miRNA size.

Overall the RNA size profile is similar to what can be observed in mouse sperm, where the dominating small RNA class is around 32 nt long(58). Mapping to the RefSeq entries confirmed reads mapping to tRNA fragments and ribosomal RNA to be the most abundant (Figure 3b). A comparison of specifically miRNAs to mouse sperm miRNA payload revealed that 3 of the 10 mostly abundant miRNAs across aged sparrows and sparrows in their prime were also among the 10 most abundant adult mouse sperm miRNAs (miR-let-7a-5p, miR-let-7f, miR-10a-5p) (Figure 3a,b). The presence of sperm miRNAs suggest a role post-fertilization, as described in mouse(7). Additionally, tgu-miR-7644, tgu-miR-2989, tgu-miR-2962 and tgu-miR-2989 was originally thought to be zebra finch specific and have not been identified in chickens(59). Identification of these miRNA demonstrate that these are shared between house sparrows and zebra finches and potentially specific to Passeriformes.

The second approach of miRNA analysis using miRDeep2 also revealed a range of miRNAs with a seed sequence shared with known zebra finch miRNAs. This could potentially implicate overlapping functionality in the regulation of mRNA targets. However, zebra finch sperm RNA has not been investigated yet, hence the comparison is restricted to somatic zebra finch miRNAs. It is also important to note that the miRDeep2 algorithm was trained and benchmarked using miRNA sequencing datasets derived mostly from somatic cells. The small RNA population in somatic cells is fundamentally different from sperm cells, due to the active fragmentation of ribosomal RNA(55) and the consequences on mRNA and ribosomal RNA abundance relative to small RNA in sperm.

The observed high abundance of tRNA fragments in sperm also suggests a role of this RNA class post fertilization reminiscent of what has been shown in mice(10, 11). In mice, in opposition to the vital role of miRNAs post-fertilization(7) tRNA fragments seem to be implicated in the transmission of effects of environmental exposures.

House sparrows hold great promise for studies on RNA based epigenetic germline inheritance since sperm can be sampled repeatedly and non-invasively. Sampling from the same individual multiple times as it ages might be a crucial refinement to not only overcome the increased variability due to the outbred background, but also to observe longitudinal effects that take place as individuals senesce. Future analysis of the contribution of miRNAs in gene regulation in sparrows will benefit strongly from this first description of sperm RNA content. Specifically in a germline context with relevance for transgenerational effects, our study paves the way for refined study design to overcome the hurdles of non-inbred animals with higher genetic complexity. Finally, our analysis has implications for ecology, as the first sparrow mirBase entries provide a new tool to study avian miRNA mediated gene regulation in a non-laboratory species.

## Acknowledgments

We thank the staff of Sanger institutes research support facility for assistance with rodent husbandry, Kay Harnish for sequencing support, Marc Ridyard and Miranda Landgraf for logistic and administrative assistance.

## Data availability

All data were deposited to and are available under accession number E-MTAB-7603 and E-MTAB-7617 at ArrayExpress (https://www.ebi.ac.uk/arrayexpress/).

## Figure legends

This manuscript contains supplementary material:

1. Supplementary Figures
2. Supplementary Tables 1-4

Supplementary Table 1

Count table representing reads mapping to known zebra finch miRNA transcripts.

Supplementary Table 2

Count table representing reads identified as miRNAs by miRDeep2 across all sparrow sperm samples indicating whether seed sequence is conserved in zebra finches.

Supplementary Table 3

(a) Results of DESeq2 differential expression analysis of aged versus prime sparrow sperm small RNA sequencing reads using zebra finch miRNA miRBase entries.

(b) Results of DESeq2 differential expression analysis of aged versus prime sparrow sperm small RNA sequencing reads using miRDeep2.

Supplementary Tabel 4

(a) Read counts of all tRNA fragments across all sparrow sperm small RNA libraries.

(b) Results of DESeq2 differential expression analysis of aged versus prime sparrow sperm small RNA sequencing reads using zebra finch tRNA annotation.

