## Supplementary material for "Mature sperm small RNA profile in the sparrow: implications for transgenerational effects of age on fitness"

Supplementary Figure 1a

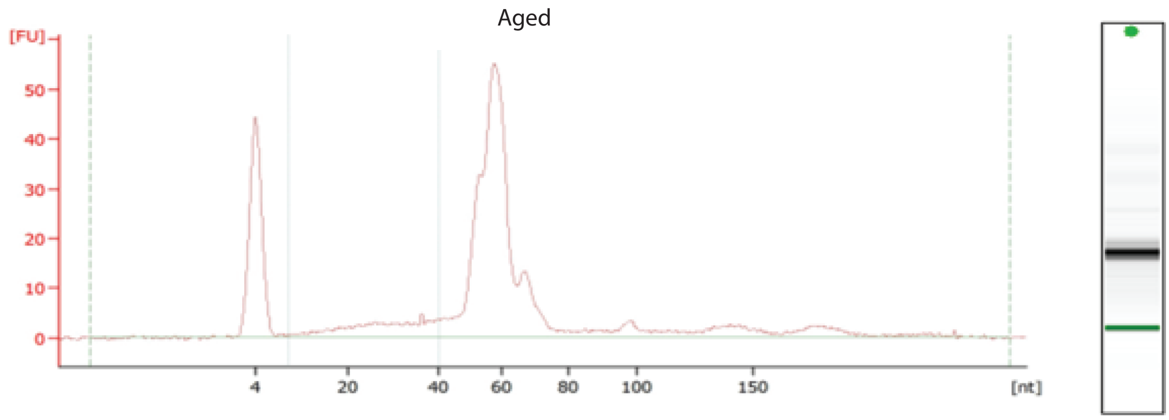


Exemplary electropherogram of sperm RNA isolateded from sparrows as obtained by the Bioanalyzer small RNA Kit.

Supplementary Figure 1b

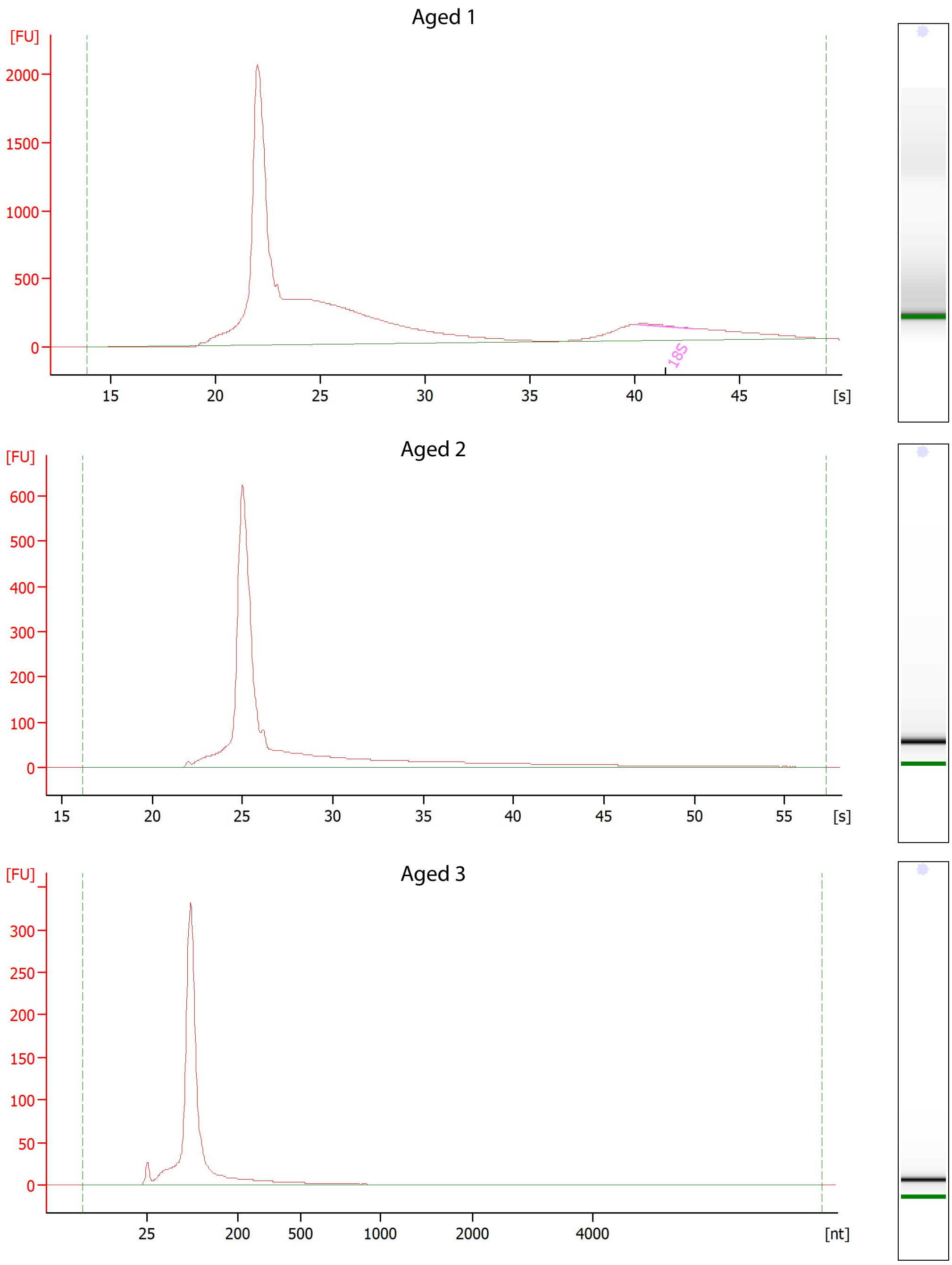

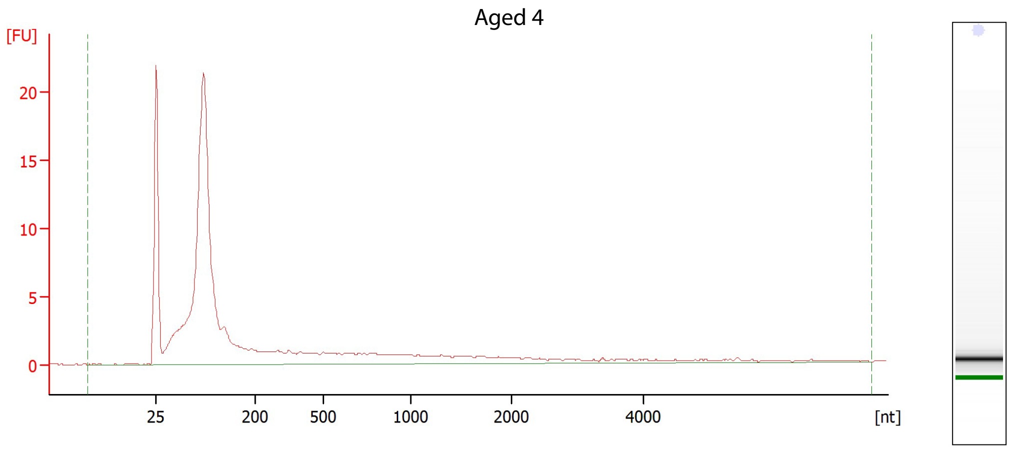
Aged 4
[FU]
20
15
10
5
0
25
200
500
1000
2000
4000
[nt]

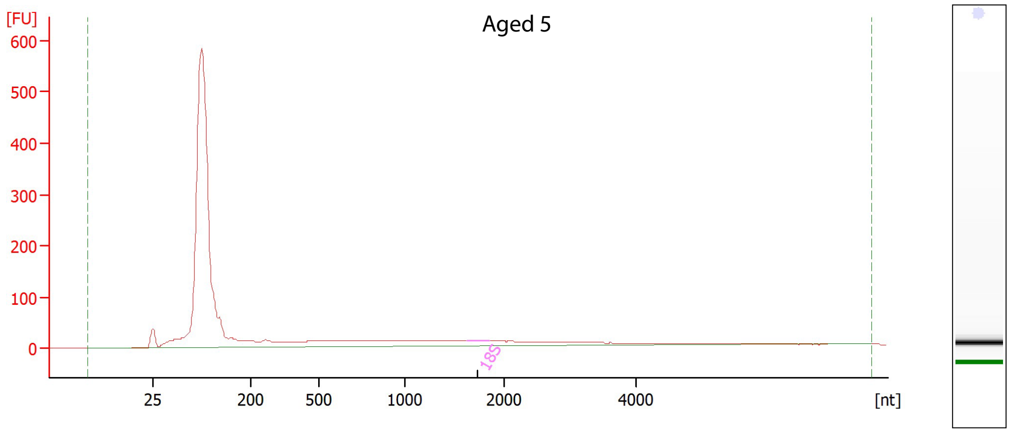
Aged 5
[FU]
600
500
400
300
200
100
0
18S
25
200
500
1000
2000
4000
[nt]

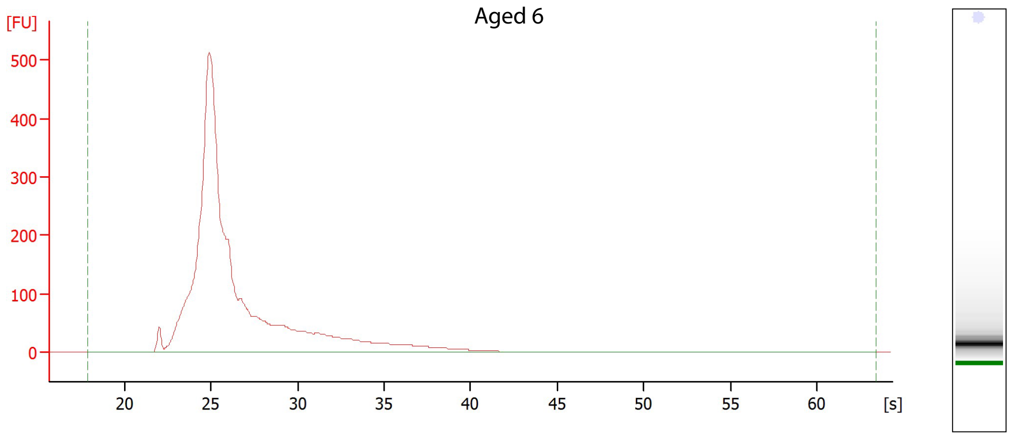
Aged 6
[FU]
500
400
300
200
100
0
20
25
30
35
40
45
50
55
60
[s]

### Aged 7

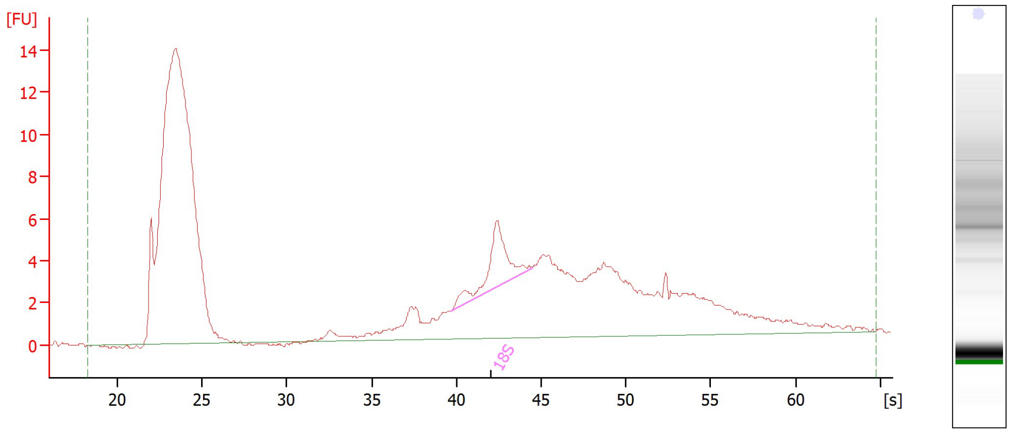


### Aged 8

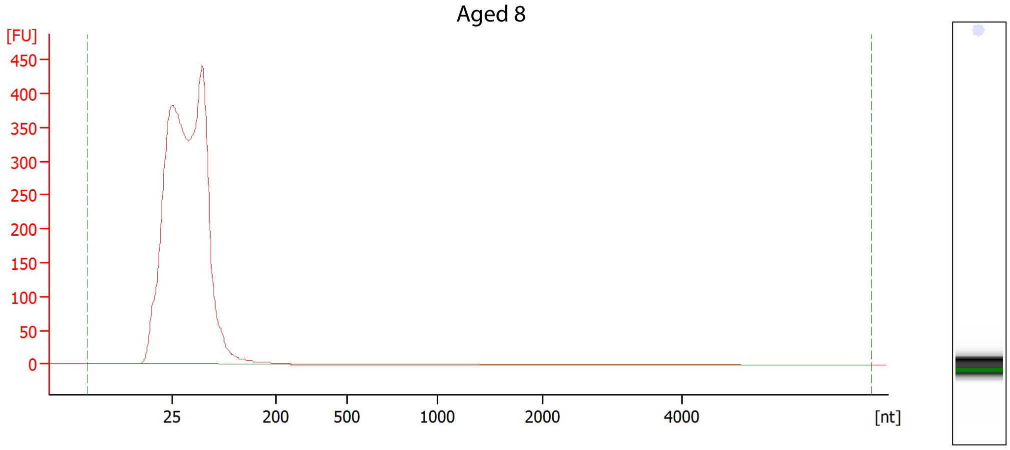


### Aged 9

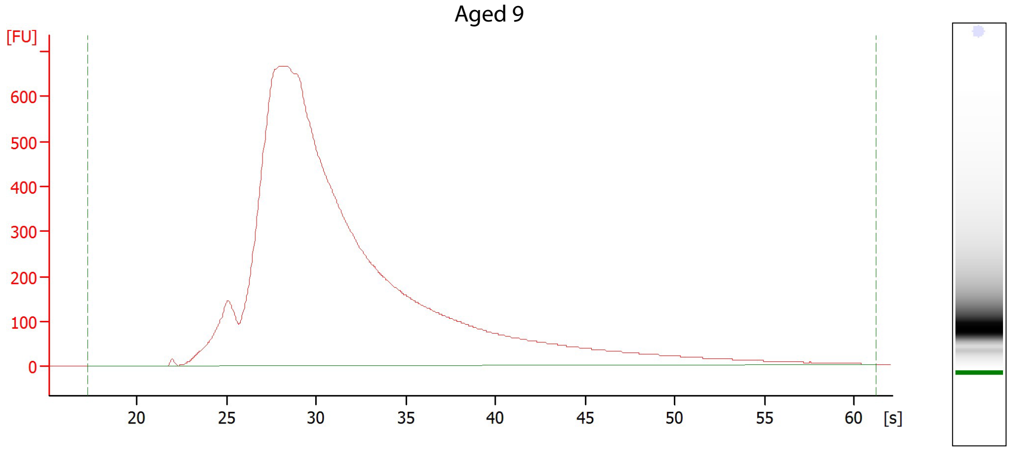

Prime 1

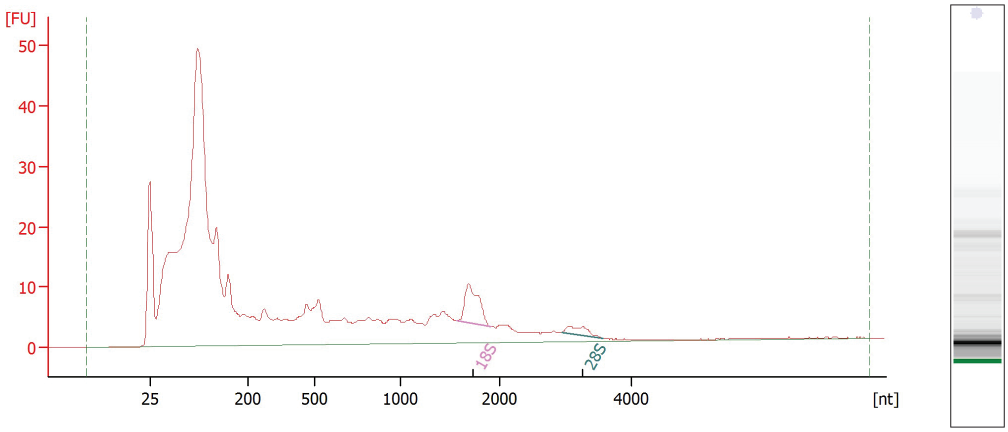


Prime 2

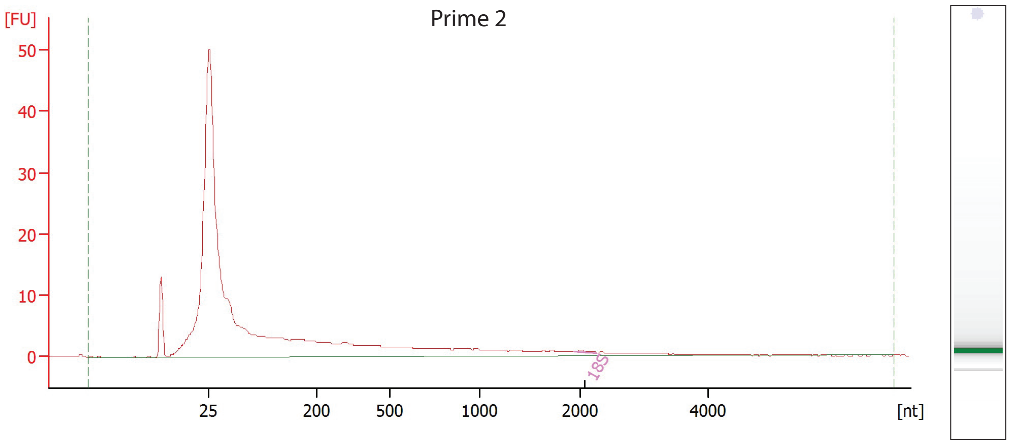


Prime 3

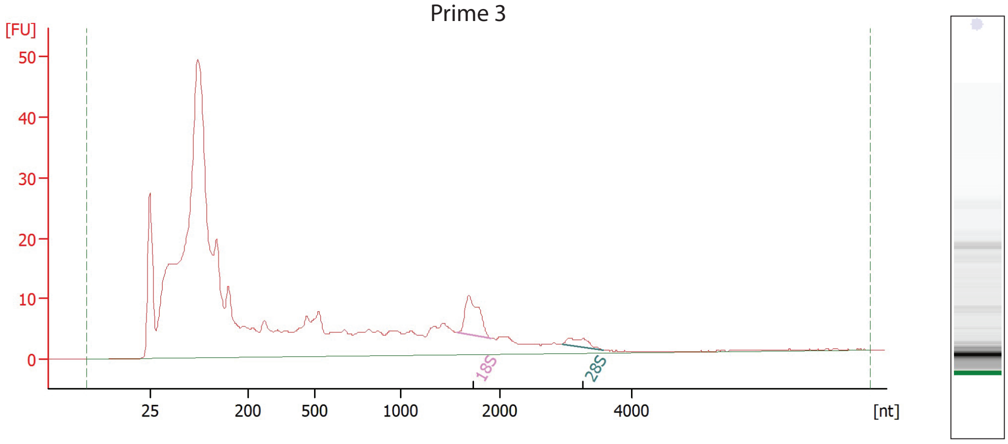

Prime 4

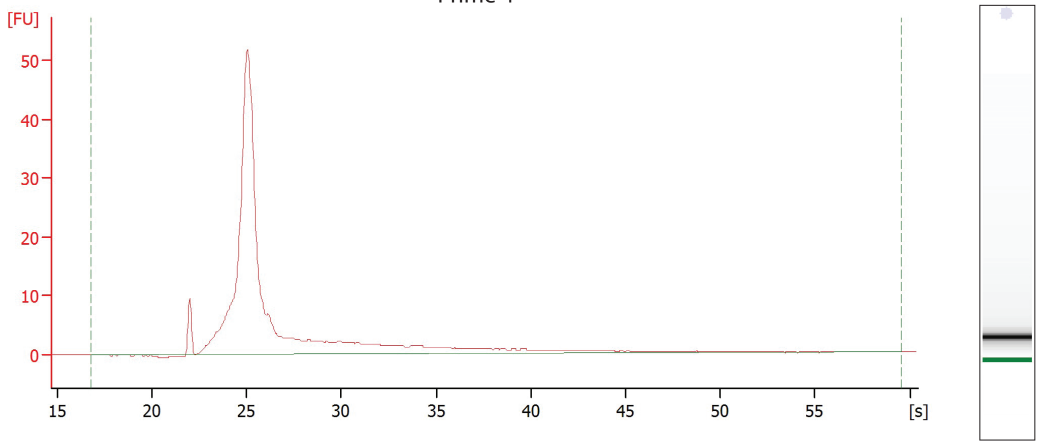


Prime 5

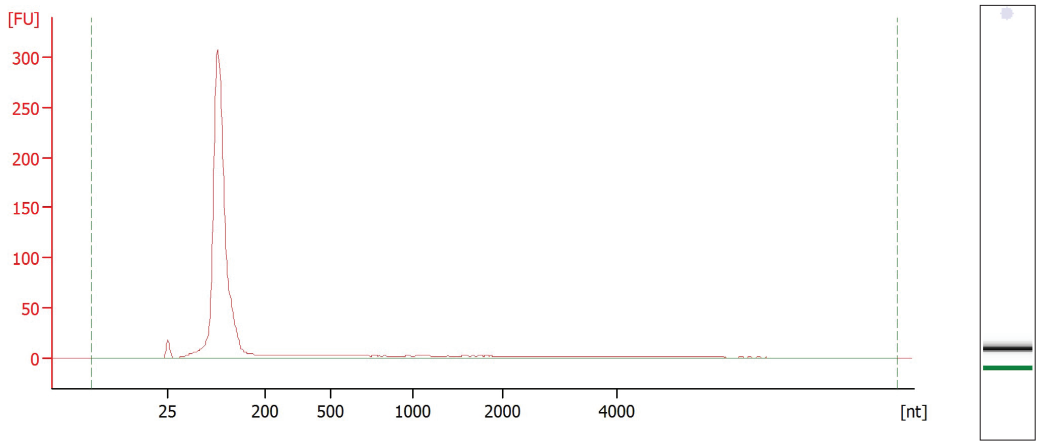


Prime 6

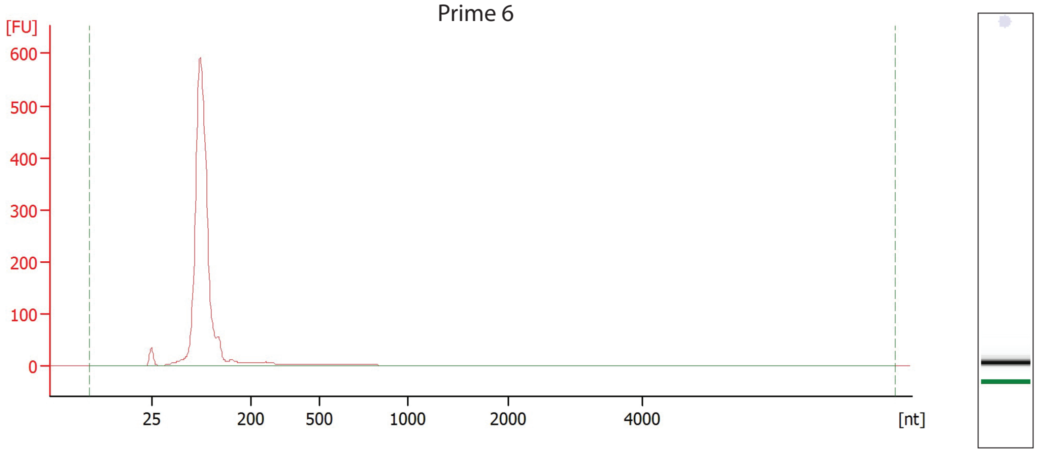

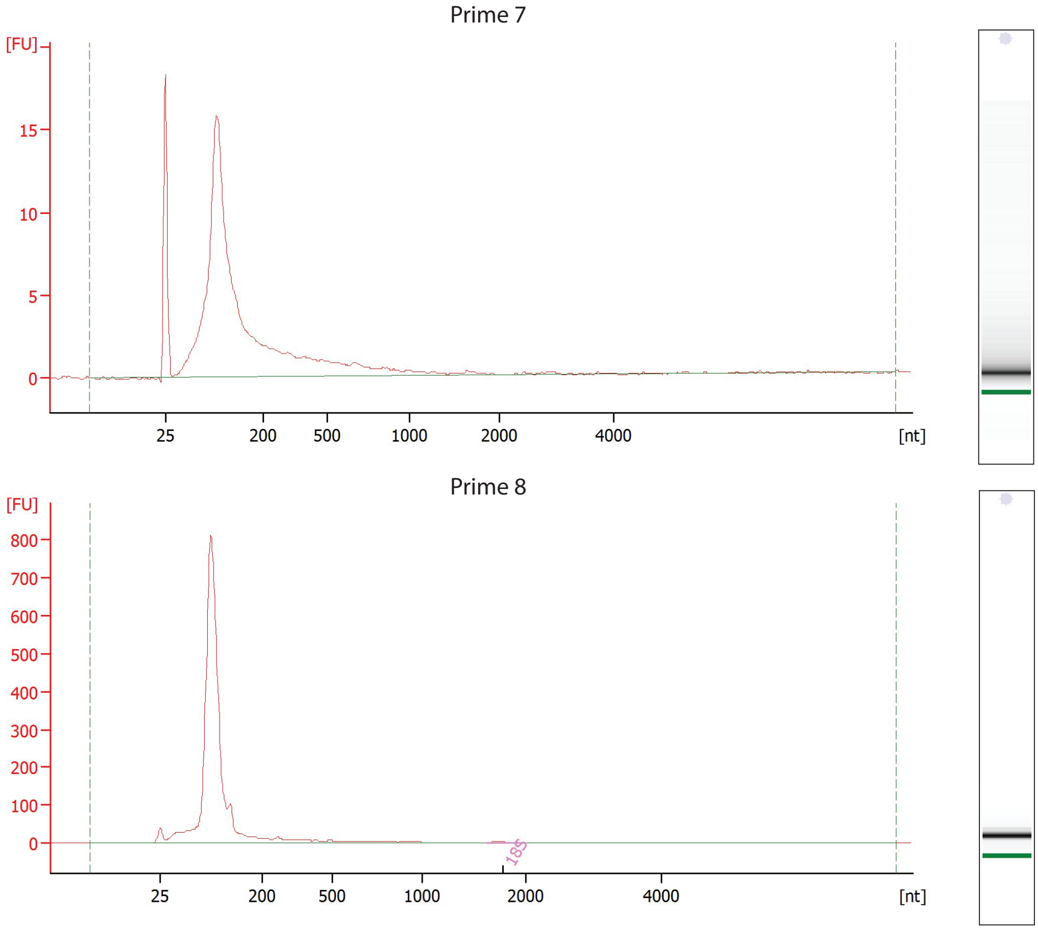


Electropherograms of sperm RNA isolateded from aged and sparrows in their prime as obtained by the Bioanalyzer Pico 6000 Kit.
Y-axis: 200 nucleotides (nt) correspond to 28 seconds (s).
